# Detecting directional epistasis and dominance from cross-line analyses in alpine populations of *Arabidopsis thaliana*

**DOI:** 10.1101/2023.04.19.537438

**Authors:** Arnaud Le Rouzic, Marie Roumet, Alex Widmer, Josselin Clo

## Abstract

The contribution of non-additive genetic effects to the genetic architecture of fitness, and to the evolutionary potential of populations, has been a topic of theoretical and empirical interest for a long time. Yet, the empirical study of these effects in natural populations remains scarce, perhaps because measuring dominance and epistasis relies heavily on experimental line crosses. In this study, we explored the contribution of dominance and epistasis in natural alpine populations of *Arabidopsis thaliana*, for two fitness traits, the dry biomass and the estimated number of siliques, measured in a greenhouse. We found that, on average, crosses between inbred lines of *A. thaliana* led to mid-parent heterosis for dry biomass, but outbreeding depression for estimated number of siliques. While heterosis for dry biomass was due to dominance, we found that outbreeding depression for estimated number of siliques could be attributed to the breakdown of beneficial epistatic interactions. We simulated and discussed the implication of these results for the adaptive potential of the studied populations, as well as the use of line-cross analyses to detect non-additive genetic effects.

## INTRODUCTION

The evolutionary forces underlying divergence between populations, in particular the relative strength of genetic drift compared to natural selection, can be revealed by studying the consequences of hybridization (Lynch, 1991; Demuth and Wade, 2005; Fitzpatrick, 2008). For example, when the evolution of populations has been mainly driven by genetic drift (e.g. due to demographic events or mating systems, Barrett *et al*., 2014), populations have likely fixed deleterious mutations (Kimura *et al*., 1963). Hybrids between populations that have accumulated (partially) recessive deleterious mutations are thus expected to display heterosis, *i.e.* an increase in fitness in F1 hybrids compared to the average fitness of their parents (Crow, 1948; Lynch, 1991; Glémin, 2003). When selection is at least as strong than drift, divergent populations are expected to accumulate genetic incompatibilities and hybrids may perform poorly compared to the parental populations, in F1 and/or subsequent generations (from the F2), a phenomenon called outbreeding depression (Lynch, 1991). Several non-exclusive genetic architectures of fitness can lead to outbreeding depression: chromosomal rearrangements, which leads to the production of aneuploid gametes in heterozygotes (Lande, 1985; Charlesworth, 1992; Kirkpatrick and Barton, 2006); under-dominance, which leads to lower fitness for heterozygotes compared to the homozygote genotypes (Schierup and Christiansen, 1996); and negative epistatic interactions among divergent alleles that are brought together for the first time in hybrids (Dobzhansky, 1937; Lynch, 1991; Demuth and Wade, 2005).

Estimating non-additive genetic effects is important because they are expected to modify how a population responds to selection (Kelly, 1999; Carter *et al*., 2005). Epistasis has the potential to modify the short- and long-term adaptive potential of a species (Cheverud & Routman, 1995; Carter *et al*., 2005; Hansen, 2015). Positive epistasis (i.e. epistatic interactions that increase traits’ values) tends to amplify the genetic variance of the trait, and consequently, the capacity of the population to respond to selection will also increase (Carter *et al*., 2005). Negative epistasis (i.e. epistatic interactions that decrease traits’ values) will have the opposite effect. Despite its potential effect on response to selection, estimates of epistasis for morphological traits are currently rare and not consistent (Le Rouzic, 2014). Pavlicev *et al*. (2010) found that epistasis tends to decrease the value of body-composition traits in *Mus musculus*, while in several plant species, epistasis tend to increase, on average, the value of different floral morphological traits (Johansen-Morris & Latta, 2006; Monnahan & Kelly, 2015; Oakley *et al*., 2015; Clo *et al*., 2021). Epistasis for fitness components has received substantially more interest. Theoretical work has indeed repeatedly pointed out that epistasis underlying fitness should drive many diversity-generating mechanisms, including the evolution of sex, recombination, and mutation rates (Phillips *et al*., 2000). Yet, empirical estimates don’t show a clear directionality (Maisnier-Patin *et al*., 2005; Kouyos *et al*., 2007; Bakerlee *et al*., 2022).

Dominance in quantitative genetics received less theoretical attention because its consequences on the evolutionary potential of a species are complex (Walsh & Lynch, 2018). Dominance can modify the adaptive potential of a trait (Clo *et al*., 2019, Clo & Opedal 2021, Sztepanacz *et al*., 2023). However, its effect depends on inbreeding (Kelly, 1999): dominance can only contribute to the covariance between parents and offspring in inbred populations, which does not occur under random mating (Falconer, 1996; Lynch & Walsh, 1998). The adaptive potential is thus not only described by the additive variance (Cockerham & Weir, 1984; Wright & Cockerham, 1985), as dominance contributes to the evolvability of quantitative traits in inbred populations (Clo & Opedal, 2021). Dominance is routinely observed for fitness components, as shown by the ubiquity of inbreeding depression (Charlesworth & Willis, 2009), but less is known about morphological traits, for which dominance effects are highly variable and can either increase or decrease traits’ values (Shaw *et al*., 1998; Kelly & Arathi, 2003; Oakley *et al*., 2015; Clo *et al*., 2021).

Several experimental protocols have been proposed to detect these non-additive genetic effects (see Le Rouzic, 2014 for a list of methods to detect epistasis). Among all, line-cross analyses have been the most used (Demuth & Wade, 2005), probably due to the simplicity of the crossing design and associated statistics. This method requires the organisms to be grown and measured in controlled conditions (generally laboratories for animals, or greenhouses for plants) to minimize the environmental variance and maximize the statistical power to detect genetic effects (Walsh & Lynch, 2018).

Predominantly selfing species are of particular interest when studying the phenotypic and fitness consequences of hybridization. By organizing populations in a mosaic of repeated fully homozygous genetic lines (Siol *et al*., 2008; Volis *et al*., 2010; Jullien *et al*., 2019), self-fertilization simplifies the dissection of the genetic architecture of hybrid fitness. Indeed, experimental hybridization between inbred lines produces F1 hybrids that are expected to be fully heterozygous at all sites where the parents differ. At each subsequent generation of selfing, the heterozygosity is divided by two, allowing one to decipher the relative contribution of dominance and epistasis to hybrids’ performance (Demuth and Wade, 2005; Fitzpatrick, 2008).

*Arabidopsis thaliana* is a natural choice to study the genetic architecture of fitness and the consequences of non-additive genetic effects in plants. *A. thaliana* (L.) Heyhn. (Brassicaceae) is native to Eurasia and North Africa but is now widely distributed throughout the Northern hemisphere (Hoffmann, 2002). This species occurs in diverse environments, and has been reported along a wide altitudinal range, from sea level up to 2000 m in the central Alps (Hoffmann, 2002). Unlike close relatives, *A. thaliana* is a predominantly self-fertilizing, annual species; average outcrossing rates in natural populations have been reported to vary between 0.3% and 2.5% (Abbott & Gomes, 1989; Bergelson *et al*., 1998; Picó *et al*., 2008).

In this study, we explored the genetic architecture of two fitness traits in natural Alpine populations of *A. thaliana*: the dry biomass and the estimated number of siliques. We first found that, on average, crosses between inbred lines of *A. thaliana* led to heterosis for dry biomass, but outbreeding depression for the estimated number of siliques. We found that heterosis for dry biomass could be attributed to a positive effect of dominance. This is likely due to the masking of recessive deleterious mutations segregating in the different inbred lines. For the estimated number of siliques, however, we found that outbreeding depression was likely due epistatic interactions. We simulated the response to selection of our traits and found that both dominance and epistasis can potentially affect the response to selection compared to additive scenarios.

## MATERIAL AND METHODS

### Study populations

We studied six natural alpine populations located along an altitudinal gradient in the Alps in the Saas Valley (Valais, Switzerland). Focal populations (Table S1) were selected from those studied by Luo *et al*. (2015). Three populations are from low altitudes (*i.e.* altitudes ranging from 850 to 1000m) and three from close to the high-elevational range margin of the species in the Alps (*i.e.* altitudes ranging from 1792 to 2012m). Distances among populations ranged from 0.8 to 25.8 km, with average distances of 6.3 km among low-altitude and 1.9 km among high-altitude populations. For our crossing experiment, we used offspring of plants collected in 2007 that were propagated in the greenhouse for three generations by single-seed descent to standardize maternal effects.

### Experimental design and measured traits

Seeds of plants originally collected in 2007 and propagated for three generations by single seed descent in a greenhouse were used as parental lines in the crossing experiment. From each of the six study populations we randomly selected four parental lines with different genotypes based on results reported in Luo *et al*. (2015). Briefly, Luo *et al*. (2015) genotyped individuals using twenty-two microsatellite markers (Table S2); the populations’ structure was reported in their Table 1 and Table S1. In the following section, when an analysis is referring to be done at the “lines” scale, it means we compared results between parental lines (seeds propagated for three generations by single seed descents). When we are referring to analyses done at the “populations” scale, we compared results between populations. All parental lines were bred to produce four offspring categories: (i) spontaneous self-fertilization, leading to the same parental genotype, (ii) outcrossing with pollen from another parental line from the same local population, (iii) outcrossing with a parental line from another population of the same altitude, and (iv) outcrossing with a parental line from another population of the other altitude. For each parental line, self-fertilization and the three cross types were realized on four different plants. Crosses were performed using the pollen of plants exclusively grown for pollen production. Parental lines were randomly matched without replacement such that all four parental lines of each population were included as both seed and pollen parents in all cross types (Table S3). The F2 generation was produced by the spontaneous self-fertilization of one individual from each F1 family (for an overview of the crossing design see Table S3). Crosses were realized by emasculating and manually pollinating 4 to 6 flowers per female/mother plant. To avoid uncontrolled cross-pollination between neighboring plants, unused flowers were removed, and pots were individually packed in Arabisifter floral sleeves (Lehle Seeds, Round Rock, Texas, USA). To ensure self-fertilization and limit accidental crossing events, selfing plants were also packed in floral sleeves during the flowering stage. No control emasculations were performed.

**Table 1.**
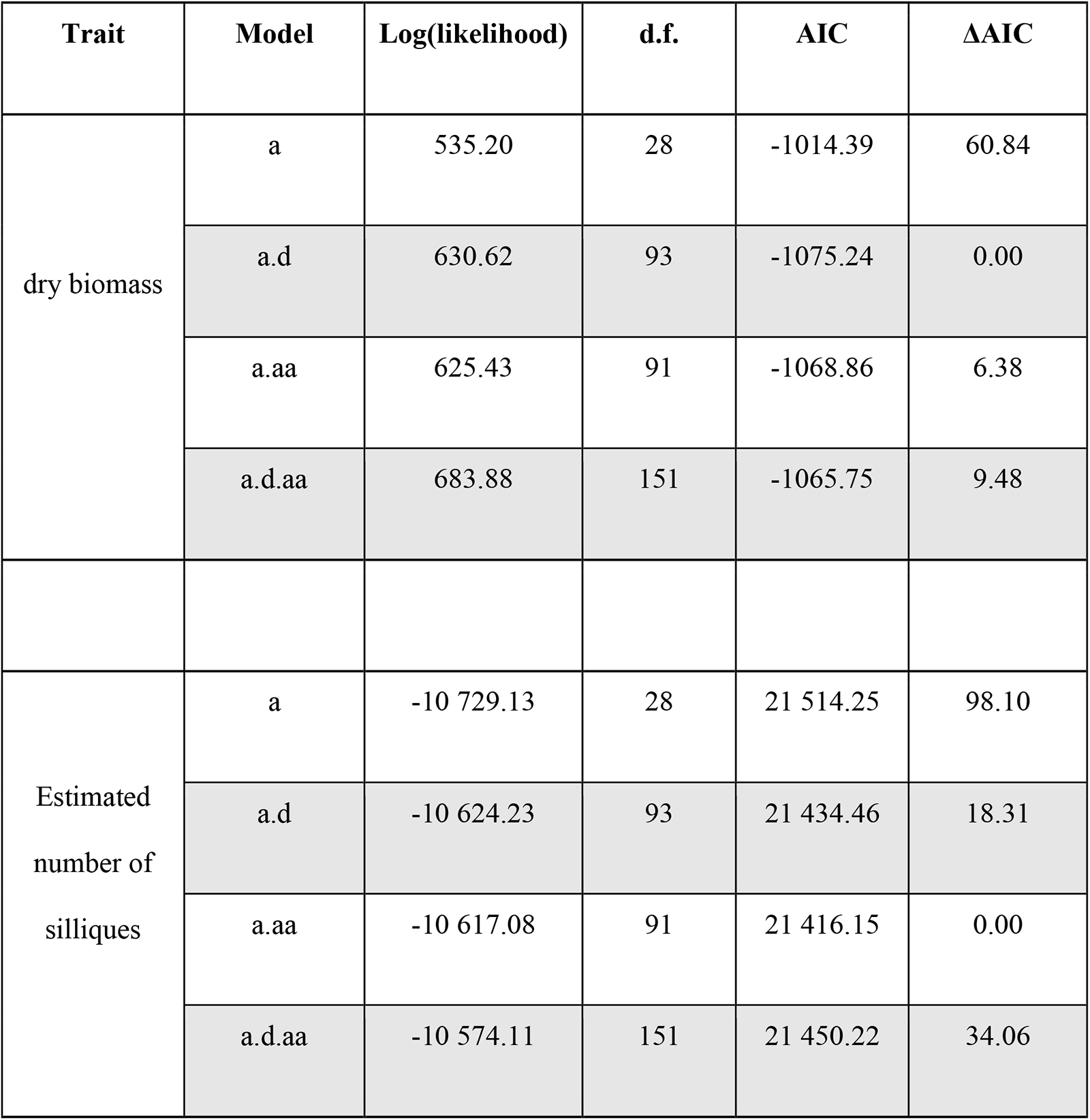
Summary of the statistical models fitted to data, when analyses are performed at the scale of genetic lines, for dry biomass and estimated number of siliques. In the table, “a” stands for additive, “d” for dominance, and “aa” for additive-by-additive epistasis. ΔAIC is the difference in AIC values between the observed and best models, differences of 2 AIC units or more are generally considered as solid statistical support for the best model.

| Trait | Model | Log(likelihood) | d.f. | AIC | ΔAIC |
| --- | --- | --- | --- | --- | --- |
| dry biomass | a | 535.20 | 28 | -1014.39 | 60.84 |
|  | a.d | 630.62 | 93 | -1075.24 | 0.00 |
|  | a.aa | 625.43 | 91 | -1068.86 | 6.38 |
|  | a.d.aa | 683.88 | 151 | -1065.75 | 9.48 |
| Estimated number of siliques | a | -10 729.13 | 28 | 21 514.25 | 98.10 |
|  | a.d | -10 624.23 | 93 | 21 434.46 | 18.31 |
|  | a.aa | -10 617.08 | 91 | 21 416.15 | 0.00 |
|  | a.d.aa | -10 574.11 | 151 | 21 450.22 | 34.06 |

Performance and phenotypic variation in all F1 and F2 families were assessed in a single large greenhouse experiment conducted in spring 2014. Seeds were sown on March 4th 2014 and stratified at 4 °C in the dark for six days. Plants were then grown in a greenhouse at ETH Zurich research station Lindau-Eschikon under long-day conditions (*i.e.* 10^4^ Lux light for 16 h, dark for 8 h; 22 °C/18 °C day/night temperatures). From each parental line, we grew 12 parental plants, 24 F1 and 48 F2 offspring. The F1 included six offspring derived from selfing and six offspring derived from each of the three cross types. The F2 generation encompassed six selfed offspring and 14 offspring from each of the three cross types. In total, the experiment encompassed 1728 plants in total (for details see Table S3).

Plants were grown individually in 7✕7✕8 cm pots randomly arranged in two greenhouse compartments, and were filled with Biouniversalerde (Oekohum GmbH, Herrenhof, Switzerland), an all-purpose soil without peat. Within each greenhouse compartment, pots were randomly arranged in 24-pot-trays. To avoid edge effects, trays were placed on tables next to each other and surrounded by “border plants” (*i.e.* plants derived by self-pollination from the study populations, sown and grown under the same conditions as the experimental plants). Trays with experimental plants were randomized twice a week until the maturation of siliques. All plants were harvested on July 1st, 2014, approximately four months after germination, when all plants that reached the flowering stage started to dry. Plants were first dried for 48h at 45°C.

We then measured the dry biomass and estimated the number of siliques per plant. To estimate the number of siliques, we first separated the different branches and isolated the reproductive sections of all branches (*i.e.* the parts of the branches carrying fruits); second, we weighted the reproductive (’reproductive weight’) sections of all branches of each individual together to the nearest 0.0001g using a Mettler AE 240 analytical balance. Third, we assessed the estimated number of siliques along three randomly selected and weighted reproductive sections and estimated the number of siliques per gram (‘silique density’); fourth, we estimated the total number of siliques produced per plant (’silique number’) as the product of the ‘silique density’ and the ‘reproductive weight’. This last measurement, the ‘silique number’, was used as a proxy for individual fecundity.

### Genetic model

Traditional line cross models consider only two parental lines and generally define genetic effects by measuring the difference between parental lines and F1 to the F2 generations. Using the F2 generations as a reference generally simplifies the mathematical expressions (Lynch & Walsh, 1998). As we aim here to analyze several line crosses at once, we reparametrized this model by taking the grand mean of all the parental lines µ as a reference (Table S4, Figure S1). The average phenotypic means of P_i_ and P_j_, as well as their intercrosses F_1, ij_ and F_2, ij_ can be expressed as:

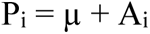

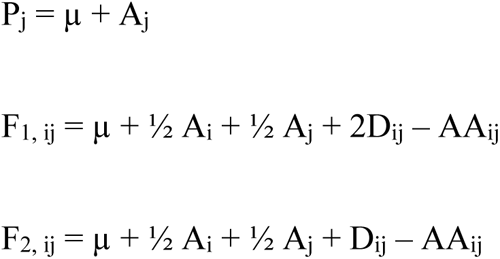

This setting defines one additive effect A per parent, and as many dominance (D) and additive-by-additive (AA) epistasis parameters as independent crosses. In the absence of backcrosses, additive-by-dominance epistatic effects cannot be identified and are merged with additive effects. Dominance-by-dominance interactions, as well as higher-order epistatic terms, had to be ignored. Note that positive dominance tends to generate heterosis (F1 and F2 generations larger than the mid-parent), while positive epistasis tends to generate outbreeding depression (F1 and F2 lower than the mid-parent).

Line-cross models aim to measure individual deviations from additivity, and were thus analyzed with a fixed-effect linear model:

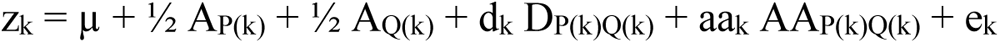

for individual k of phenotype z_k_, of parents from parents P(k) and Q(k), with d_k_ = aa_k_ = 0 if P(k) = Q(k), aa_k_ = -1 if P(k) ≠ Q(k) (k is from an intercross F_1_ or F_2_), and d_k_ = 2 (or =1) if k results from an F_1_ (or an F_2_) intercross. e_k_ is a Gaussian-distributed residual of variance V_e_.

Four models of various complexity were fit to each dataset: Additive (only the additive terms A_i_ were considered), Dominance (A_i_ and D_ij_), Epistasis (A_i_ and AA_ij_), and Full model (A_i_, D_ij_, and AA_ij_). The four models were compared by a model selection procedure based on the Akaike Information Criterion (AIC, Anderson & Burnham, 2004) ; the best model had the lowest AIC value; AIC differences larger than 2 units were considered to be a significantly poorer fit to the data. Models were fit independently on dry mass and estimated number of siliques, and both genetic differentiation levels (Lines and Population) were considered.

### Simulated adaptive trajectories

We used the best model and the associated non-additive genetic effects to simulate the consequences of non-additivity and inbreeding on the response to selection for the traits of interest, compared to an additive model. In a two-locus, two-allele context, we assumed infinite populations and linkage equilibrium. Noting p_1_ the frequency of allele A1 (1-p_1_ the frequency of allele B1) at locus 1, and s the selfing rate, the genotype frequencies at locus 1 deviate from the Hardy-Weinberg equilibrium (Hartl & Clark 1989) :

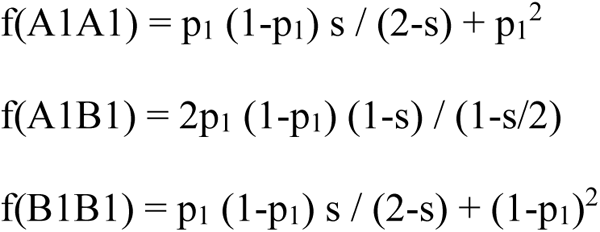

The genotype frequencies at locus 2 followed the same logic (with p_2_ the frequency of allele A2 and 1-p_2_ the frequency of allele B2). Assuming linkage equilibrium, the frequencies of double genotypes were computed as the product of single genotype frequencies. Genotype-phenotype maps were parameterized according to the most supported model (additive effects and dominance for weight, additive and AA epistasis for siliques), and parameterized according to a traditional F2 model:

|  | Locus 1 |  |  |
| --- | --- | --- | --- |
| Locus 2 | A1A1 | A1B1 | B1B1 |
| A2A2 | $\mu - a - d + aa$ | $\mu - a/2$ | $\mu - aa + d$ |
| A2B2 | $\mu - a/2$ | $\mu + d$ | $\mu + a/2$ |
| B2B2 | $\mu - aa - d$ | $\mu + a/2$ | $\mu + a - d + aa$ |

When non-zero, parameter values were the average of pairwise effects (red bars in Fig 2): for weight, µ = 0.43g, a=0.11g, d=0.03g; for siliques: µ=779, a=262, aa=83. In this setting, µ stands for the mean random-mating F2 population. Fitness was proportional to the phenotype; the lowest fitness genotype was 80% that of the best (i.e., fitnesses ranged between 0.8 and 1). Genotype frequencies were recomputed and normalized after selection (e.g., f’(A1A1) = f(A1A1) w_A1A1_ / w, where w stands for the mean fitness), and allele frequencies after selection (e.g. p_1_’ = f’(A1A1) + f’(A1B1)/2) were used to compute genotype frequencies at the next generation. Starting allele frequencies were p_1_ = p_2_ = ½ at both loci; the phenotype of starting populations was not necessarily at µ, as starting populations were not F2 due to selfing. The deterministic simulation procedure was iterated for 50 generations, which was in practice enough to reach stable mean phenotypic values.

## RESULTS & DISCUSSION

We found that performing the analyses at the scale of the genetic lines or at the scale of the populations gave compatible results, although population-level analysis had lower statistical power due to the smaller sample size. As within-population variation was often of the same magnitude as between-population variation, averaging lineages within a population was not justified, and we decided to present the results for the genetic lines in the main text, the results at the population scale being available as supplementary material (Table S4, Figures S2 and S3).

### Consequences of experimental hybridization on dry biomass and the production of siliques

Our first question was related to the consequences of non-additive genetic effects on the phenotype of hybrid (F1 and F2) populations compared to their parents. The raw phenotypes of parental lines and within-population crosses are available as supplementary materials (Figure S3). We found that, on average, F1 hybrids exhibited heterosis for dry biomass (Figure 1), with an increase of 9.6% (0.454g in F1 hybrids), compared to the mean parental value of 0.414g. This amount of heterosis is in line with what is found in other populations of *A. thaliana* (Oakley *et al*., 2015), and in other predominantly selfing species (Rhode & Cruzan, 2005; Dolgin *et al*., 2007; Volis *et al*., 2011; Gimond *et al*., 2013; Clo *et al*., 2021), for different fitness proxies (dry biomass, fruits and seeds production). In contrast, F2 hybrids had similar mean dry biomass to the parents (0.403g, -2%).

**Figure 1.**
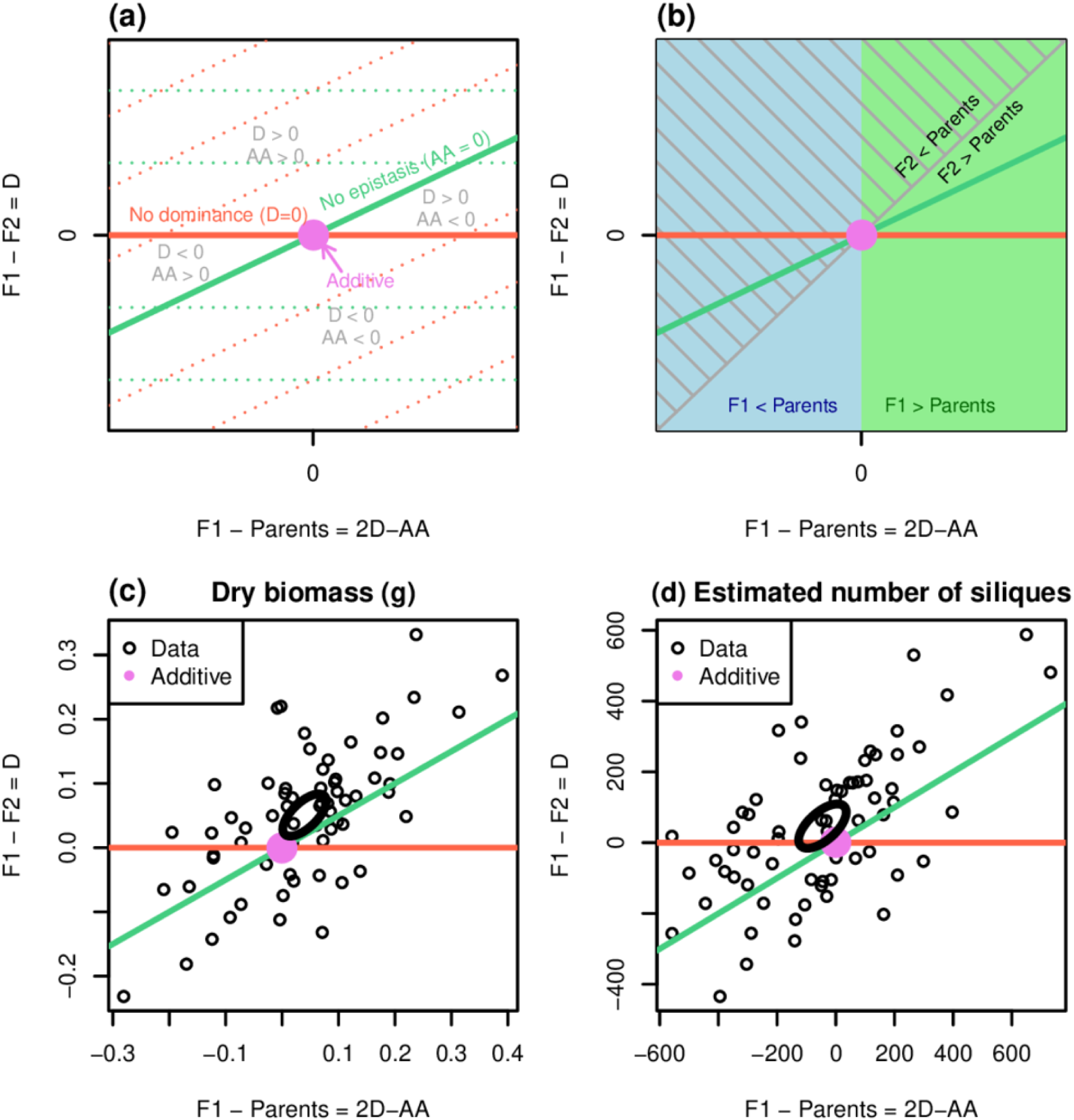
Graphical representation of the parental, F1 and F2 values, when analyses are performed at the scale of the genetic lines. **(a)** Empty representation indicating the sign of dominance and epistasis as a function of the position of the data points. The X-axis corresponds to the difference between the F1 and the mid-parent (a mixture of dominance and epistasis), while the Y-axis stands for the difference between the F2 and the F1 crosses (dominance only). **(b)** Location of heterosis and outbreeding depression for F1 (colors) and F2 (hatching) populations, with the same axes as in (a). **(c)** Distribution of data for dry biomass. **(d)** Distribution of data for estimated number of siliques. The ellipse represents the 95% confidence ellipse of the average across all crosses.

On the other hand, F1 and F2 hybrids exhibited outbreeding depression for the estimated number of siliques (Figure 1), with respectively a decrease of 5.5% and 11.3% in F1 and F2 hybrids, compared to the mean parental value. This is slightly lower than other values found for different fitness proxies (seeds and fruits production, germination rate etc.) in other predominantly selfing species (Rhode & Cruzan, 2005; Dolgin *et al*., 2007; Volis *et al*., 2011; Gimond *et al*., 2013; Oakley *et al*., 2015; Clo *et al*., 2021; Soto *et al*., 2023).

These opposite patterns for dry biomass and fruit number could be considered a surprising result because biomass is generally positively correlated with fitness components (see Younginger *et al*., 2017 for a review). However, such an observation is not unheard of. Studies in natural and laboratory accessions of *Arabidopsis thaliana* also found heterosis patterns for the dry mass and outbreeding depression for a fitness proxy (pollen viability in Nasrallah *et al*., 2000; seed production in Barth *et al*., 2003; fruit production Vasseur *et al*., 2019). In the sister species *A. lyrata,* Li *et al*. (2019) also found that selfing populations exhibit heterosis for above- and below-ground biomass, and a slight outbreeding depression pattern for fitness (measured as the probability of bolting) in outcrossed progeny of selfing populations. Finally, Clo *et al*. (2021) found that in the predominantly selfing species *Medicago truncatula,* experimental hybridization between inbred lines leads to heterosis for dry mass but outbreeding depression for seed production. It is known that environmental factors, such as nutrients or temperature, have a key role in the transition from vegetative to reproductive stages, like in flowering probability (see Cho et al. 2017 for a review). It is thus possible that the ecological conditions found in the greenhouse, which are more favorable than those in the field, might have modified trade-offs between vegetative growth and investment in reproduction; extrapolating our greenhouse results to natural populations thus relies on the assumption that environmental conditions do not affect the relative performance of genotypes (limited G✕E interactions).

We also found that, on average, plants from low-altitude populations or hybrids generated with lines from low-altitude populations were generally heavier and produced more siliques compared to high-altitude plants (Figure S4). However, this result is mainly due to a few inbred lines within each high-altitude population that performed badly, while others were in lines with what is found in low-altitude populations (Figure S3). This altitude effect might be explained by the difference in ecological conditions between low- and high-altitude populations, with for example mean annual temperature and precipitation being very different (see Luo *et al*., 2015 for details).

### Non-additive effects in natural populations of plants

We found that non-additive effects contribute to the genetic architecture of both traits. For dry mass, we found that the best model explaining the data was the one including additive and dominant genetic effects (Table 1), and the observed pattern of heterosis was mostly due to positive dominant effect (d=+0.03g on average, to be compared with an average additive effect of a=0.11g, Figure 2). Oakley *et al*. (2015) found similar results in crosses between south European and Scandinavian lineages of *A. thaliana*. The positive dominance likely reflects the positive effects of masking deleterious mutations fixed at different loci in the different selfing lines (Charlesworth 2018). The contribution of epistasis to dry biomass cannot totally be ruled out when the data are analyzed at the population level (Table S4). It indicates that both the masking of deleterious mutations and a potential synergistic effect of masking at different sites could explain the heterosis pattern (Oakley *et al*., 2015; Charlesworth 2018).

**Figure 2.**
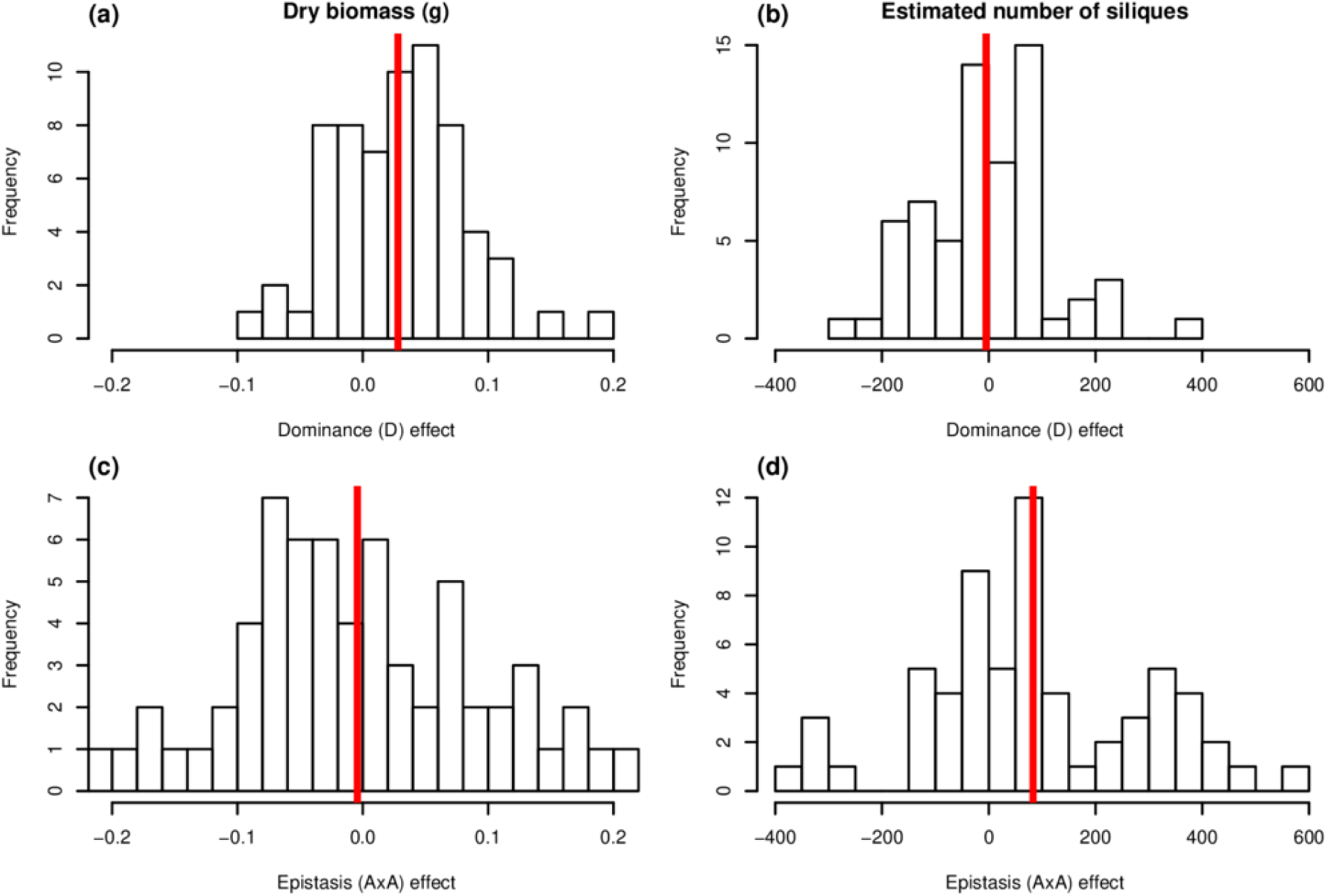
Distribution of the dominance (**(a)** and **(b)**) and epistatic (**(c)** and **(d)**) genetic effects when analyses are performed at the scale of the genetic lines, for dry biomass (**(a)** and **(c)**) and estimated number of siliques (**(b)** and **(d)**). The red lines indicate mean values.

For the estimated number of siliques, we found that the best model explaining the data was the one including additive and additive-by-additive epistatic genetic effects (Table 1), and the observed pattern of outbreeding depression was due to positive additive-by-additive epistatic interactions (average aa = 83 siliques, to be compared to an average additive effect a = 262 siliques, Figure 2). The outbreeding depression can be explained by the breakdown of positive additive-by-additive epistatic interactions found in the parental selfing lines during experimental hybridization events, as found in other species (Rhode & Cruzan, 2005; Johansen-Morris & Latta, 2006; Monnahan & Kelly, 2015; Oakley *et al*., 2015; Clo *et al*., 2021).

Finding substantial non-additive effects in fitness-related traits is not unexpected, as selection is expected to erode the additive genetic variance, exposing the non-additive variation (Roff & Emerson, 2006, Burch *et al*., 2024). Yet, dominance (dry mass) and epistasis (number of siliques) remained small compared to additive effects, suggesting that these traits might not be very correlated with fitness, or that the genetic correlation between fitness-related traits in general could be negative (so that the observed increase in dry mass or number of siliques is compensated by the decrease in unobserved fitness traits). Some remaining additive variance in fitness is also expected at mutation – selection equilibrium. Finally, strong inbreeding and/or limited gene flow could limit the local variance in fitness, while fitness differences could be revealed when artificially crossing populations.

### Implications for the adaptive potential of Alpine populations of *A. thaliana*

The distinct genetic architecture among the two fitness traits studied here implies different effects on the capacity to respond to selection. By using a two-locus and two-alleles model that mimics the genetic architecture of our traits, we explored the consequences of non-additive effects on the capacity to respond to a hypothetical selection pressure (Figure 3). When the departure to additivity is attributed to dominance, as for the dry mass, the total response to selection depends on the selfing rate (Figure 3(a)). The fact that selfing populations respond faster to selection is expected, as selfing increases the heritable variance by a factor of (1+F) due to the higher proportion of homozygotes genotypes (Wright, 1921). The fact that dominance *per se* did not affect the response to selection in selfing populations is also expected, because dominance is only expressed in heterozygote genotypes, which are rare in predominantly selfing populations/species such as *A. thaliana.* With inbreeding and dominance, the estimation of adaptative potential is different than in outcrossing species, because the whole genetic variance (in the different components of the genetic diversity) is a better predictor of the capacity to respond to selection than the additive variance (see for example Wright & Cockerham, 1985, Kelly 1999 or Clo & Opedal 2021 for further details). For estimated number of siliques, we found positive additive-by-additive epistasis. In the short term, positive epistasis tended to increase the capacity to respond to selection, as the silique production increased faster with epistasis (Figure 3(b)). This is due to the fact that positive epistasis increases the amount of additive variance of a quantitative trait, and, as a result, the capacity to respond to selection (Carter *et al*., 2005; Monnahan & Kelly, 2015). Contrary to dominance, positive epistasis is not able to change the nature of the genotype which will eventually be fixed by selection. As for dry mass, selfing populations respond faster to selection (more additive variance), but selfing does not modify the end result. In the long term, epistasis slightly increased the seed production compared to an additive model (Figure 3(b)), the difference being small because the epistatic coefficient is small compared to additive effects.

**Figure 3.**
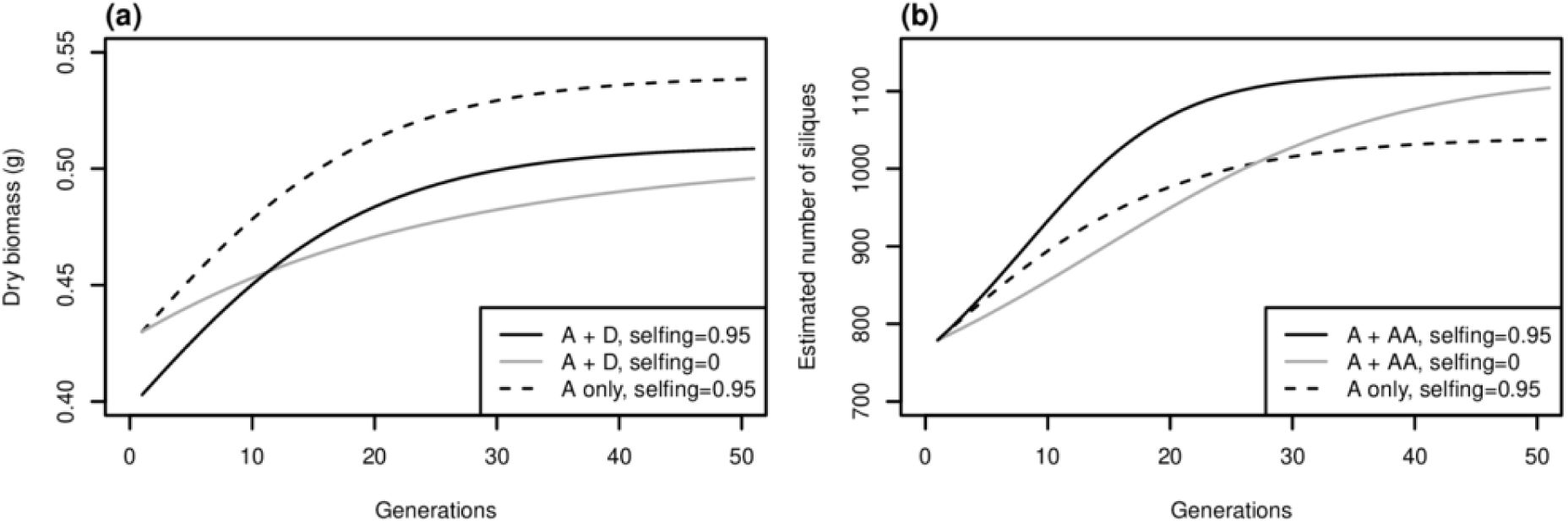
Simplified simulations of hypothetical responses to directional selection on dry biomass **(a)** and estimated number of siliques **(b)**. The additive case (dashed lines) is contrasted to two-allele two-locus genetic architectures in which the relevant non-additive effects are accounted for (dominance for dry mass, epistasis for estimated number of siliques; effects were averaged out over all pairs of populations). The selfing rate in *A. thaliana* is about 0.95; simulations with random mating populations (gray) are provided for comparison.

### Limits of the statistical model to detect the nature of non-additive effects

The line cross model we proposed makes it possible to test for the presence of dominance and epistasis, but it has potential drawbacks. First, the fact that plants have to be grown in controlled conditions makes the different traits to be very different from what is found in natural populations (see for example Weng *et al*., 2021 for estimates in natural populations of *A. thaliana*). This overestimation of the effects may make our predictive response to selection larger than expected in nature, but the pattern should be qualitatively the same if the signs of epistasis and dominance remain similar, and if the above-mentioned trade-off between vegetative growth and reproductive investment remains true in nature. Second, it can estimate epistasis, which is essential for understanding the capacity of populations to respond to selection, but it was not possible to dissect the different forms of epistatic variance (additive-by-additive, additive-by-dominant, and dominant-by-dominant). For inferring all these parameters from cross-line analyses, one needs more crosses than just the F1 and F2 individuals (see Lynch & Walsh, 1998, including reciprocal back-crosses for example, and see Oakley *et al*., 2015 for a case study). This means that the positive dominance we detected for dry biomass could be a mixture of negative dominance and epistasis, or just complex forms of epistasis (beyond additive-by-additive interaction effects). As a consequence of this identifiability problem, genetic effect estimates for dominance and epistasis tend to be statistically correlated; in other words, the model has a substantial power to detect departure from additivity, but less power to disentangle dominance and epistasis.

## CONCLUSIONS

Our study highlights the contribution of non-additive genetic effects to the genetic architecture of fitness components. Here, we found that both dominance and epistasis affect the genetic architecture of dry biomass and silique production, leading to heterosis for the dry mass and outbreeding depression for estimated number of siliques in F1 and F2 hybrids. We however found that non-additive genetic effects remain quantitatively small compared to additive components. Simulations reflecting the genetic architecture of the studied traits, as well as the mating systems of our model species *A. thaliana*, showed that both the non-additive genetic effect and the selfing rate have a significant influence on the potential to respond to selection. Testing such theoretical predictions will require artificial selection experiments (see Monnahan and Kelly, 2015; for a case study), to verify whether non-additive theoretical models of adaptation are operational in practice.

## Supplementary materials

**Table S1:**
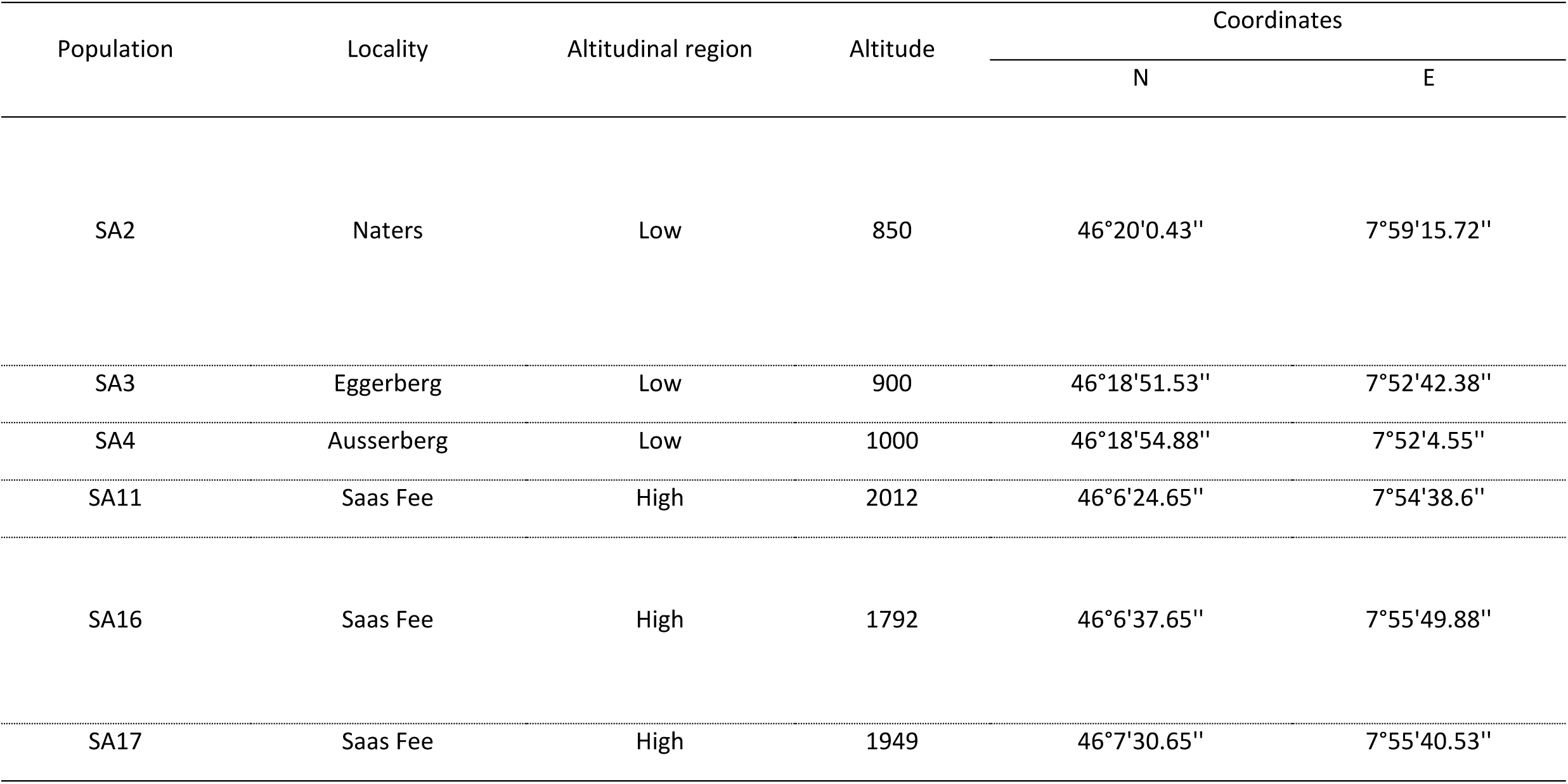
Names, locations and sampling of the studied populations.

**Table S2:**
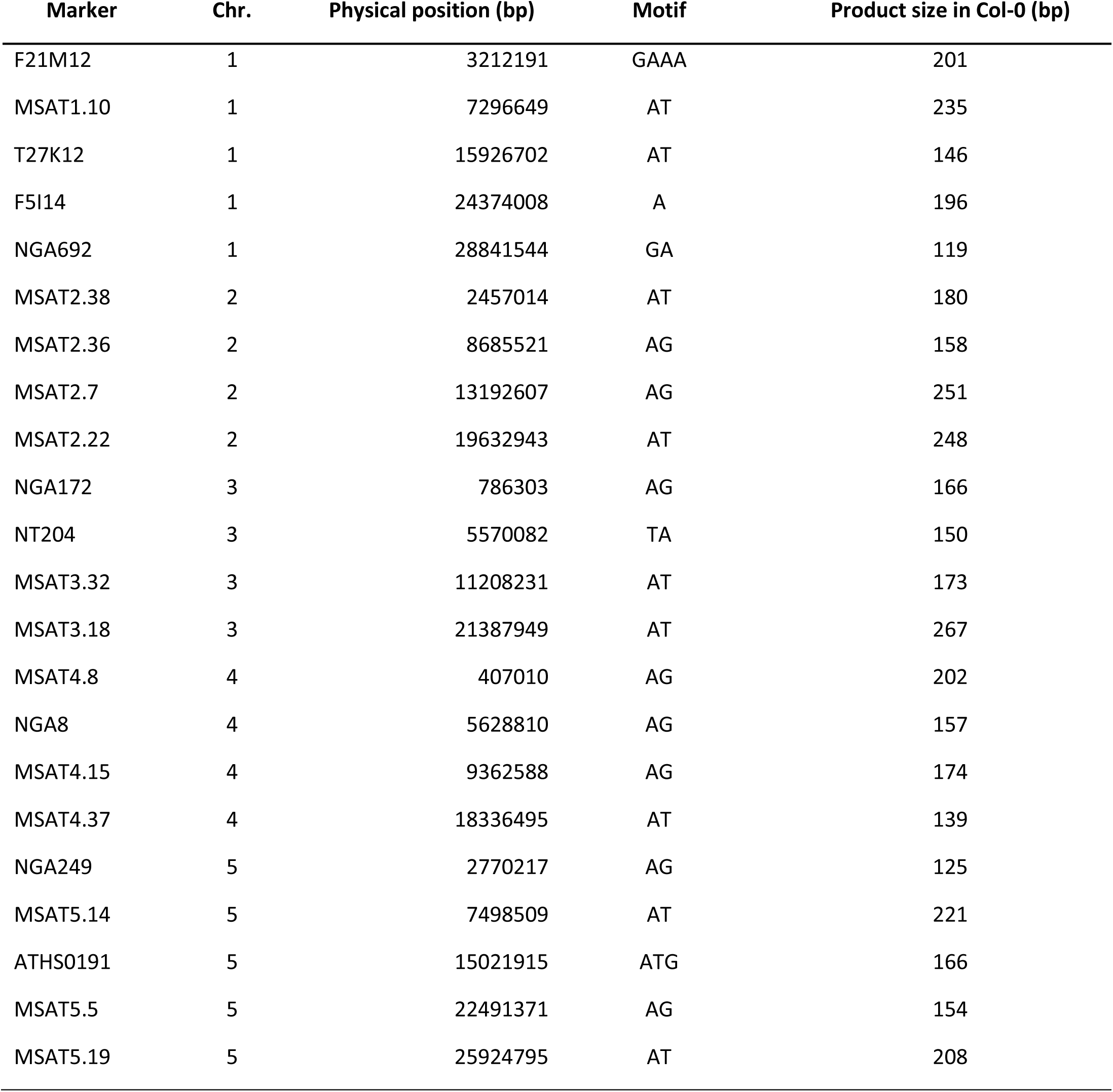
Details of microsatellite markers.

**Table S3.**
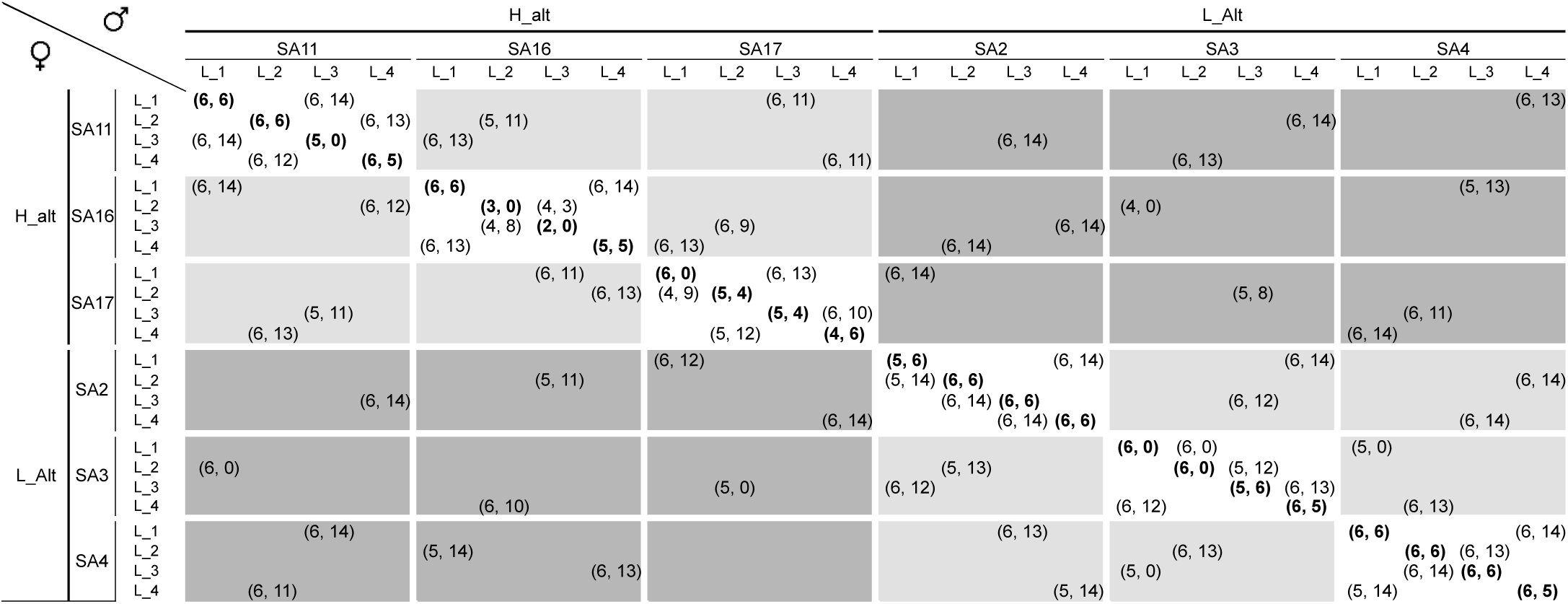
estimated number of individuals obtained through selfing (diagonal line, values in bold), and outcrossing in F1 and F2 (first and second number, respectively). Within population, between populations, and between altitudinal regions outcrosses are coloured in white, light grey, and dark grey respectively. Ideally, 6 F1 and 6 F2 selfed offspring and 6 F1 and 12 F2 outcrossed offspring were obtained. Unsuccessful crosses and/or mortality cause observed discrepancies. Abbreviations: High and low altitudinal regions: H_alt and L_alt; Parental lines: L_1, L_2, L_3, and L_4.

**Table S4.**
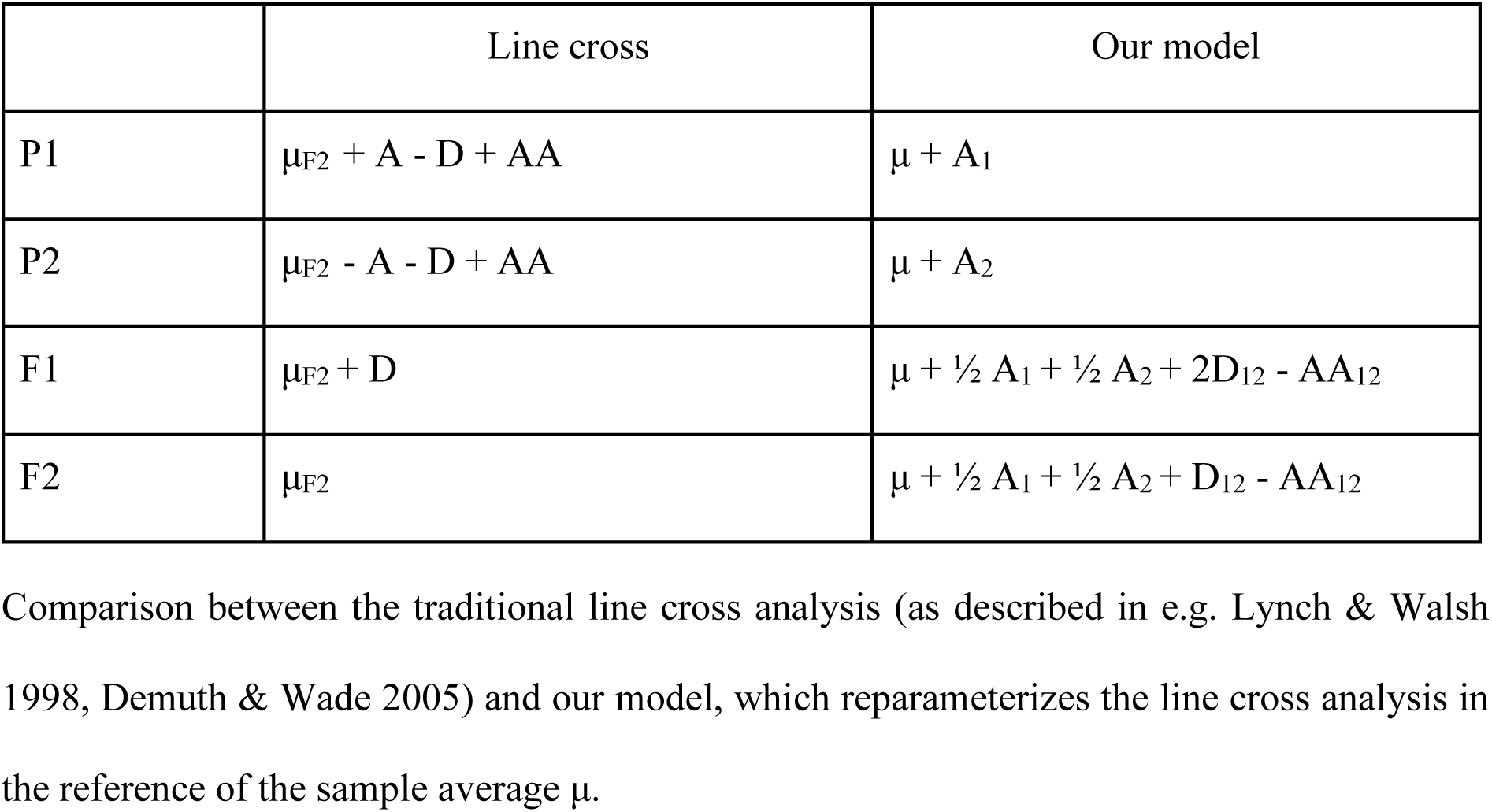

|  | Line cross | Our model |
| --- | --- | --- |
| P1 | $\mu_{F2} + A - D + AA$ | $\mu + A_1$ |
| P2 | $\mu_{F2} - A - D + AA$ | $\mu + A_2$ |
| F1 | $\mu_{F2} + D$ | $\mu + \frac{1}{2} A_1 + \frac{1}{2} A_2 + 2D_{12} - AA_{12}$ |
| F2 | $\mu_{F2}$ | $\mu + \frac{1}{2} A_1 + \frac{1}{2} A_2 + D_{12} - AA_{12}$ |

Comparison between the traditional line cross analysis (as described in e.g. Lynch & Walsh 1998, Demuth & Wade 2005) and our model, which reparameterizes the line cross analysis in the reference of the sample average $\mu$ .

**Table S5.** Summary of the statistical models fitted to data, when analyses are performed at the scale of the populations, for the dry weight and the number of silliques. ΔAIC is the difference in AIC values between the observed and best models.

| Trait | Model | Log(likelihood) | d.f. | ΔAIC |
| --- | --- | --- | --- | --- |
| Dry weight | a | 392.30 | 7 | 77.19 |
|  | a.d | 445.57 | 22 | 0.65 |
|  | a.aa | 445.90 | 22 | 0.00 |
|  | a.d.aa | 455.73 | 37 | 10.33 |
| Number of siliques | a | -10840.90 | 7 | 98.10 |
|  | a.d | -10786.40 | 22 | 18.31 |
|  | a.aa | -10783.62 | 22 | 0.00 |
|  | a.d.aa | -10777.73 | 37 | 34.06 |

**Figure S1.**
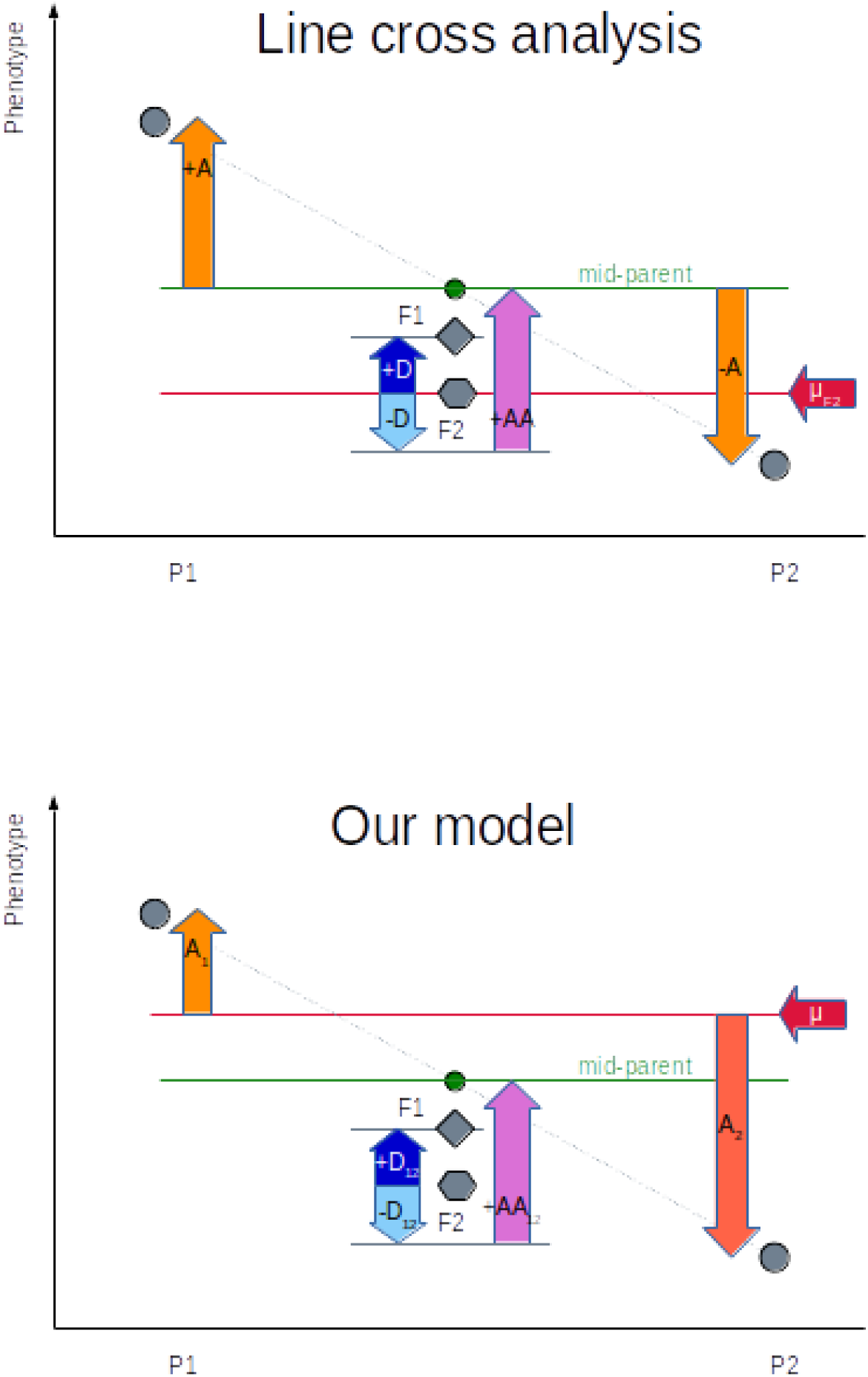
Representation of the genetic effects. Top: traditional line cross analysis between two populations; the reference (red arrow) is the F2 population, and the additive effect is measured relative to the mid-parent. Dominance is the difference between F1 and F2, and epistasis measures the difference between the F2 (from which dominance has been removed) and the mid-parent. Bottom: we changed the reference to the grand mean of the sample (μ), so that additive effects are population-specific. Dominance and epistasis are specific to a pair of populations, and keep the same meaning as in the line cross model.

**Figure S2.**
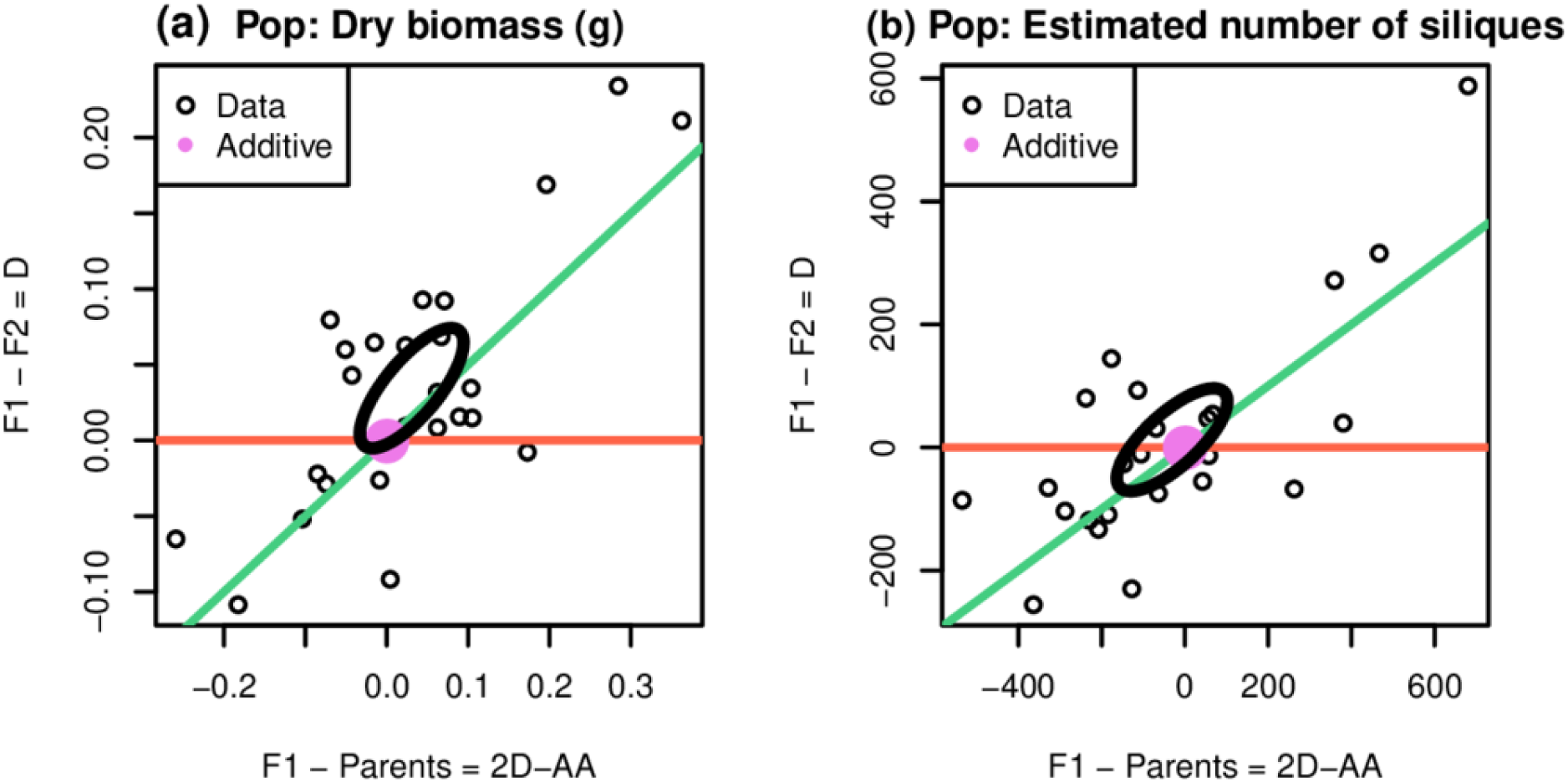
Graphical representation of the parental, *F*_1_ and *F*_2_ values, when analyses are performed at the scale of the populations. **(a)** Distribution of data for the dry biomass. **(b)** Distribution of data for the estimated number of siliques.

**Figure S3.**
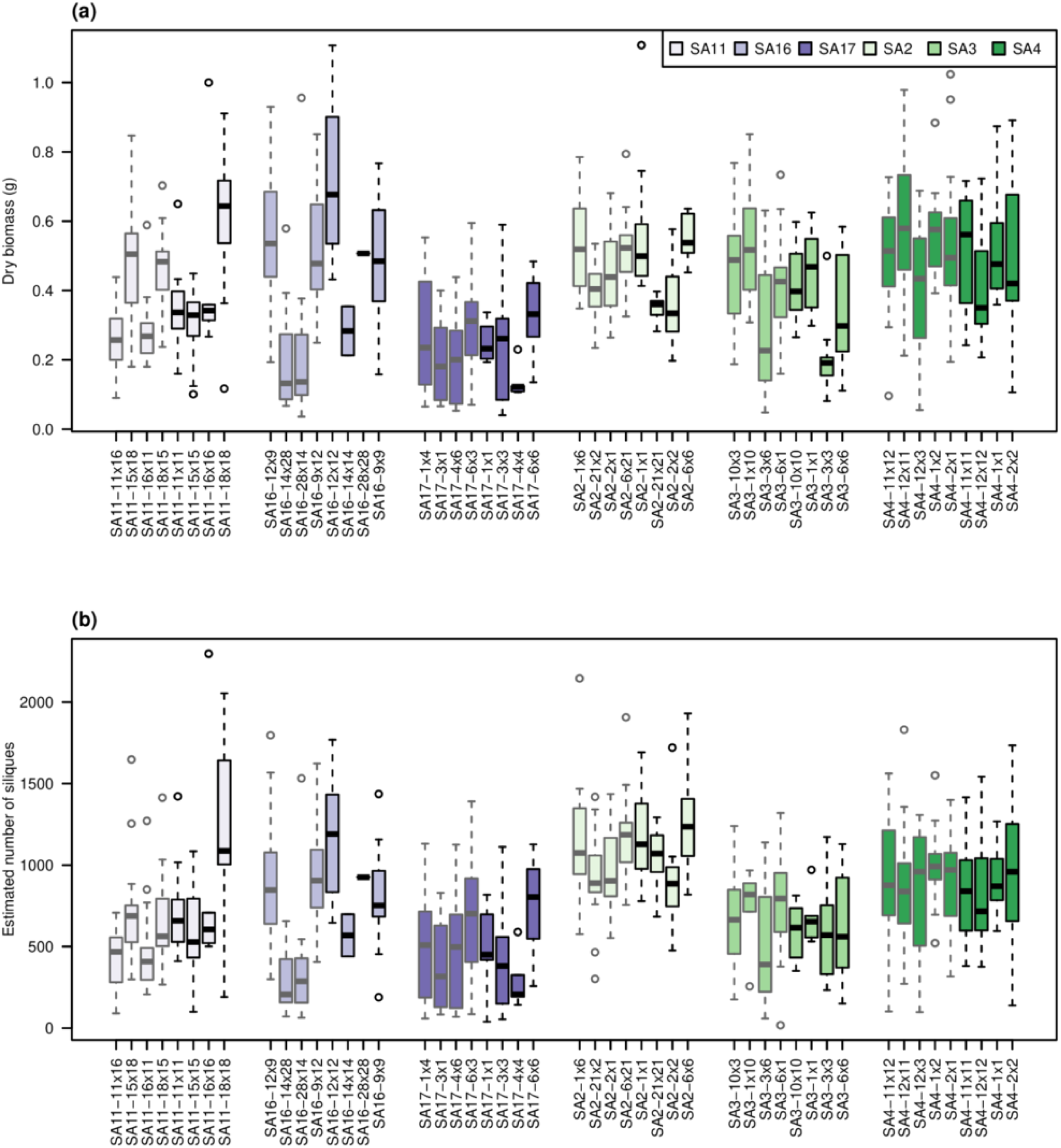
Distribution of the raw phenotypic values of within-population crosses, for the dry biomass **(a)** and the estimated number of siliques **(b)**. Colors identify maternal populations (blue plots = high altitude populations, green plots = low altitude populations); boxplots indicate the quartiles and the median of the distributions.

**Figure S4.**
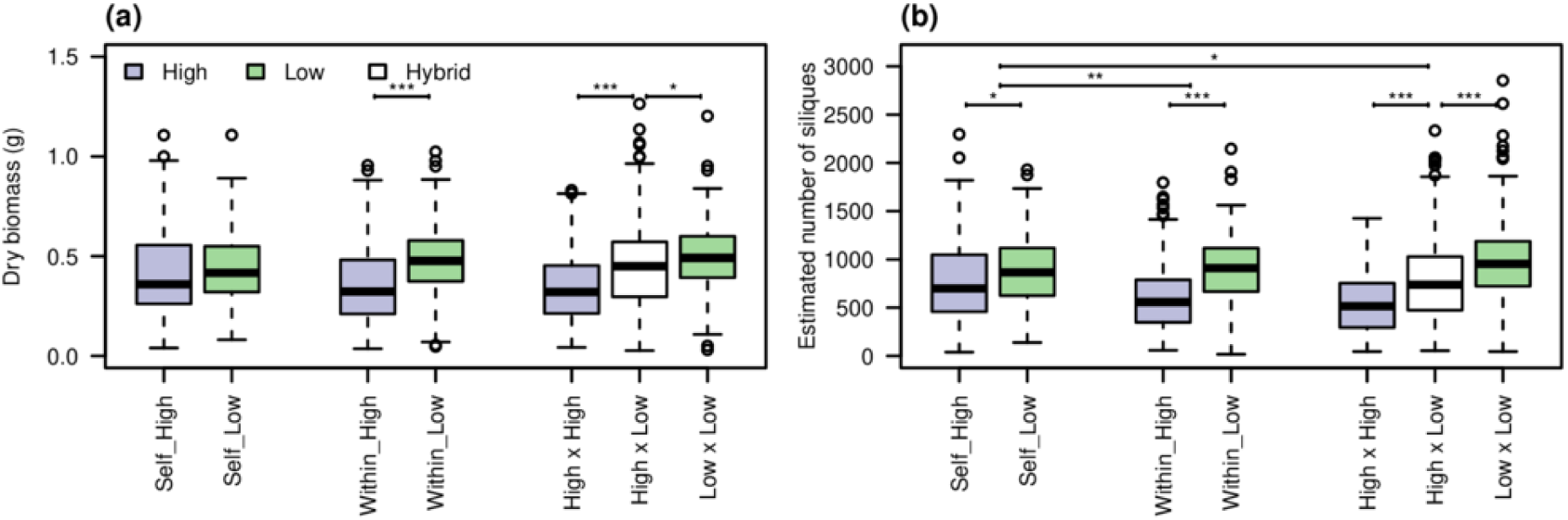
Distribution of the phenotypic values for the different kind of crosses (F1 and F2) and the altitude of populations, for the dry biomass **(a)** and the estimated number of siliques **(b)**. Boxplots indicate the quartiles and the median of the distributions, significance has been tested with *t-*tests. * = *p* < 0.05 ; ** = *p* < 0.01 : *** = *p* < 0.001

## Notes

https://github.com/lerouzic/Epicross

### Competing Interest Statement

The authors have declared no competing interest.

### Summary of Updates

We added more details about the theoretical model and corrected typos

