## Supplementary materials for "Detecting directional epistasis and dominance from cross-line analyses in alpine populations of *Arabidopsis thaliana*"

Table S1: Names, locations and sampling of the studied populations

| Population | Locality | Altitudinal region | Altitude | Coordinates | |
| --- | --- | --- | --- | --- | --- |
|  |  |  |  | N | E |
| SA2 | Naters | Low | 850 | 46°20'0.43'' | 7°59'15.72'' |
| SA3 | Eggerberg | Low | 900 | 46°18'51.53'' | 7°52'42.38'' |
| SA4 | Ausserberg | Low | 1000 | 46°18'54.88'' | 7°52'4.55'' |
| SA11 | Saas Fee | High | 2012 | 46°6'24.65'' | 7°54'38.6'' |
| SA16 | Saas Fee | High | 1792 | 46°6'37.65'' | 7°55'49.88'' |
| SA17 | Saas Fee | High | 1949 | 46°7'30.65'' | 7°55'40.53'' |

|  | **Table S2**: Details of microsatellite markers | | | | |
| --- | --- | --- | --- | --- | --- |
| **Marker** | | **Chr.** | **Physical position (bp)** | **Motif** | **Product size in Col-0 (bp)** |
| F21M12 | | 1 | 3212191 | GAAA | 201 |
| MSAT1.10 | | 1 | 7296649 | AT | 235 |
| T27K12 | | 1 | 15926702 | AT | 146 |
| F5I14 | | 1 | 24374008 | A | 196 |
| NGA692 | | 1 | 28841544 | GA | 119 |
| MSAT2.38 | | 2 | 2457014 | AT | 180 |
| MSAT2.36 | | 2 | 8685521 | AG | 158 |
| MSAT2.7 | | 2 | 13192607 | AG | 251 |
| MSAT2.22 | | 2 | 19632943 | AT | 248 |
| NGA172 | | 3 | 786303 | AG | 166 |
| NT204 | | 3 | 5570082 | TA | 150 |
| MSAT3.32 | | 3 | 11208231 | AT | 173 |
| MSAT3.18 | | 3 | 21387949 | AT | 267 |
| MSAT4.8 | | 4 | 407010 | AG | 202 |
| NGA8 | | 4 | 5628810 | AG | 157 |
| MSAT4.15 | | 4 | 9362588 | AG | 174 |
| MSAT4.37 | | 4 | 18336495 | AT | 139 |
| NGA249 | | 5 | 2770217 | AG | 125 |
| MSAT5.14 | | 5 | 7498509 | AT | 221 |
| ATHS0191 | | 5 | 15021915 | ATG | 166 |
| MSAT5.5 | | 5 | 22491371 | AG | 154 |
| MSAT5.19 | | 5 | 25924795 | AT | 208 |

**Table S3.** estimated number of individuals obtained through selfing (diagonal line, values in bold), and outcrossing in F1 and F2 (first and second number, respectively). Within population, between populations, and between altitudinal regions outcrosses are coloured in white, light grey, and dark grey respectively. Ideally, 6 F1 and 6 F2 selfed offspring and 6 F1 and 12 F2 outcrossed offspring were obtained. Unsuccessful crosses and/or mortality cause observed discrepancies.

Abbreviations: High and low altitudinal regions: H_alt and L_alt; Parental lines: L_1, L_2, L_3, and L_4.

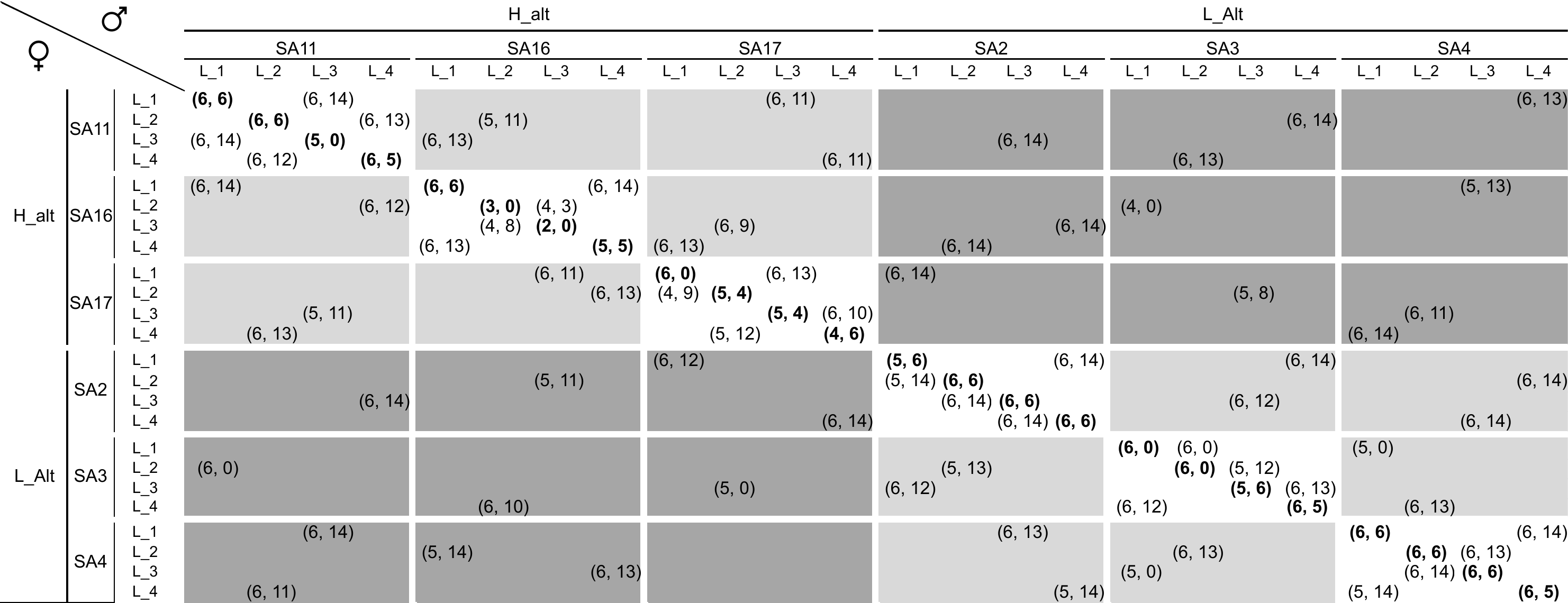

**Table S4**

|  | Line cross | Our model |
| --- | --- | --- |
| P1 | μ_F2_ + A - D + AA | μ + A_1_ |
| P2 | μ_F2_ - A - D + AA | μ + A_2_ |
| F1 | μ_F2_ + D | μ + ½ A_1_ + ½ A_2_ + 2D_12_ - AA_12_ |
| F2 | μ_F2_ | μ + ½ A_1_ + ½ A_2_ + D_12_ - AA_12_ |

Comparison between the traditional line cross analysis (as described in e.g. Lynch & Walsh 1998, Demuth & Wade 2005) and our model, which reparameterizes the line cross analysis in the reference of the sample average μ.

**Table S5.** Summary of the statistical models fitted to data, when analyses are performed at the scale of the populations, for the dry weight and the number of silliques. ΔAIC is the difference in AIC values between the observed and best models.

| **Trait** | **Model** | **Log(likelihood)** | **d.f.** | **ΔAIC** |
| --- | --- | --- | --- | --- |
| Dry weight | a | 392.30 | 7 | 77.19 |
|  | a.d | 445.57 | 22 | 0.65 |
|  | a.aa | 445.90 | 22 | 0.00 |
|  | a.d.aa | 455.73 | 37 | 10.33 |
| Number of silliques | a | -10840.90 | 7 | 98.10 |
|  | a.d | -10786.40 | 22 | 18.31 |
|  | a.aa | -10783.62 | 22 | 0.00 |
|  | a.d.aa | -10777.73 | 37 | 34.06 |

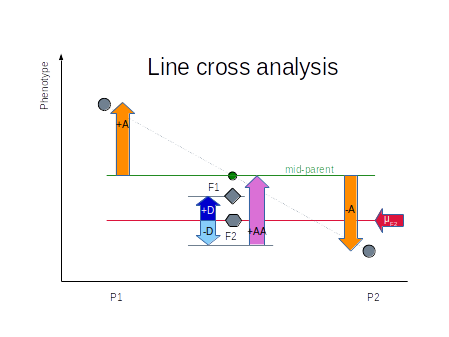

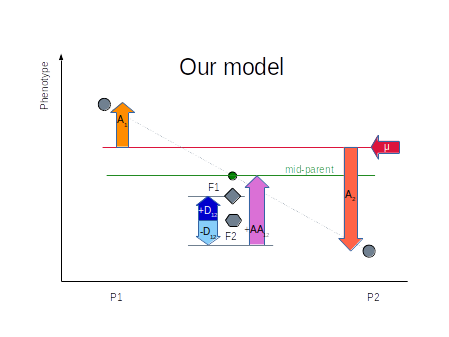

**Figure S1**. Representation of the genetic effects. Top: traditional line cross analysis between two populations; the reference (red arrow) is the F2 population, and the additive effect is measured relative to the mid-parent. Dominance is the difference between F1 and F2, and epistasis measures the difference between the F2 (from which dominance has been removed) and the mid-parent. Bottom: we changed the reference to the grand mean of the sample (μ), so that additive effects are population-specific. Dominance and epistasis are specific to a pair of populations, and keep the same meaning as in the line cross model.

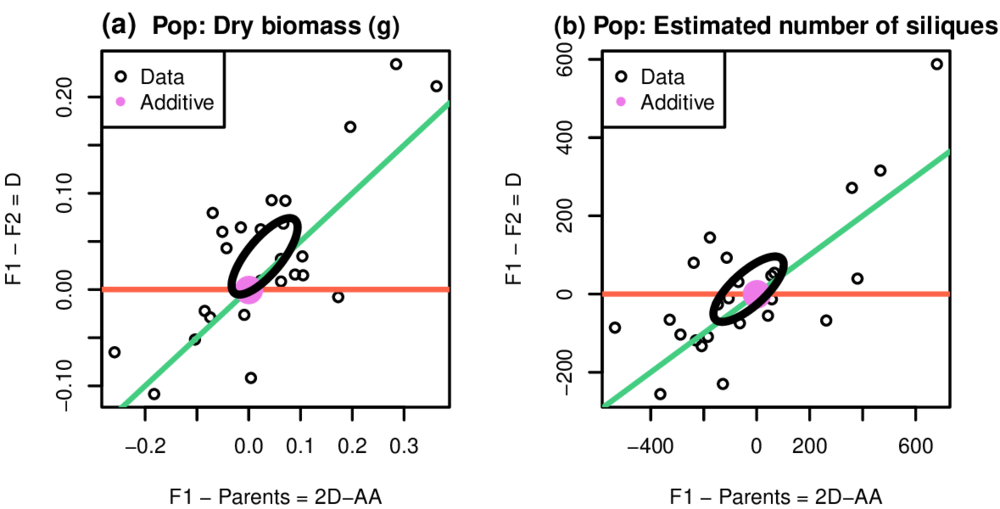

**Figure S2.** Graphical representation of the parental, *F*_1_ and *F*_2_ values, when analyses are performed at the scale of the populations. **(a)** Distribution of data for the dry biomass. **(b)** Distribution of data for the estimated number of siliques.

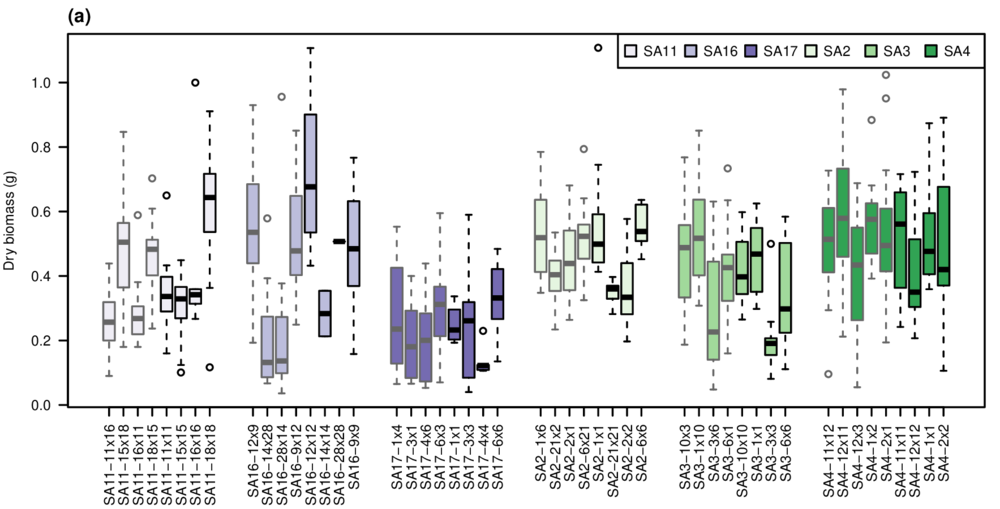

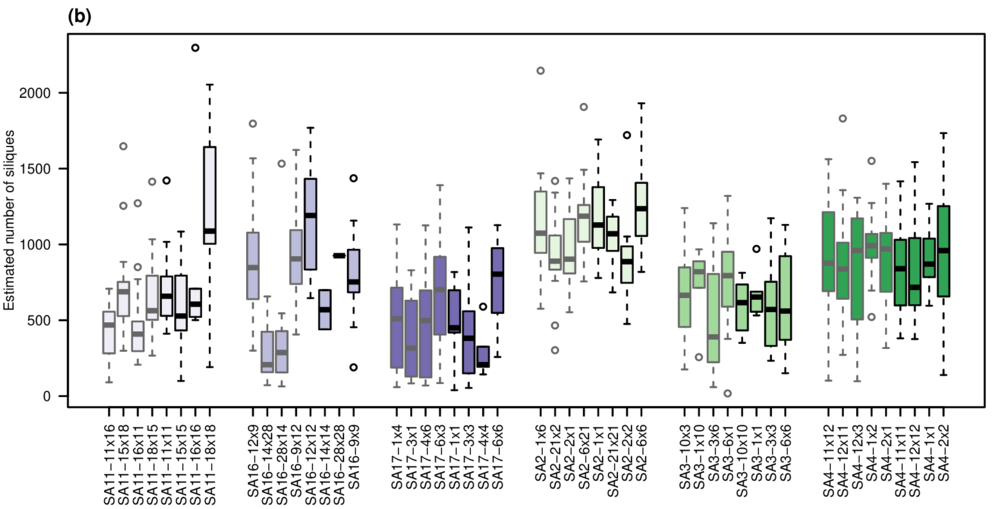

**Figure S3.** Distribution of the raw phenotypic values of within-population crosses, for the dry biomass **(a)** and the estimated number of siliques **(b)**. Colors identify maternal populations (blue plots = high altitude populations, green plots = low altitude populations); boxplots indicate the quartiles and the median of the distributions.

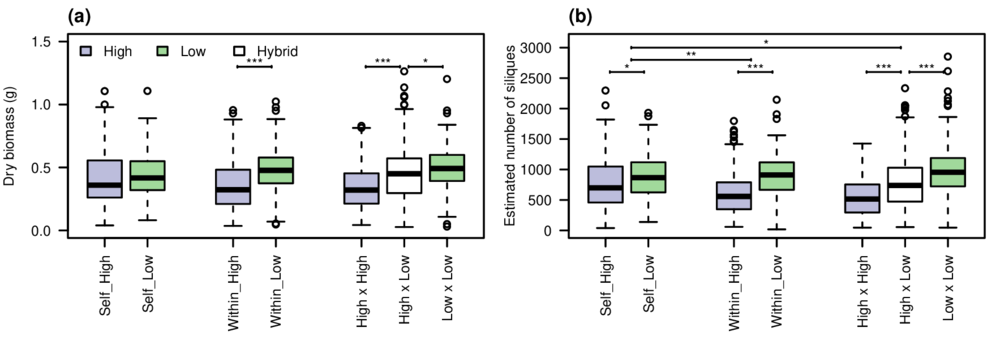

**Figure S4.** Distribution of the phenotypic values for the different kind of crosses (F1 and F2) and the altitude of populations, for the dry biomass **(a)** and the estimated number of siliques **(b)**. Boxplots indicate the quartiles and the median of the distributions, significance has been tested with *t-*tests. * = *p* < 0.05 ; ** = *p* < 0.01 : *** = *p* < 0.001
